# A platform for automated training of mammalian cell physiology

**DOI:** 10.64898/2026.08.13.744473

**Authors:** Patrick Erickson, Douglas Hazel, Ramses Martinez, Kostyantyn Shcherbina, Susan L. Marquez, Thomas Ferrante, Katarina Johnson, Angelina Pimkina, Hananel Hazan, Juanita Mathews, Adama Marie Sesay, Michael Levin

## Abstract

Controlling cell physiology is difficult, not only because of cells’ complexity, but also their capacity for real-time adaptation to interventions, leading to challenges such as drug resistance and transgene silencing. Accumulating evidence suggests that this adaptivity resembles classical forms of learning defined in behavioral science. However, a lack of appropriate platforms has led to gaps in our understanding of cells’ capacity for adaptive problem-solving in physiological and transcriptional space. Here, we present a device, the Cell Trainer, capable of performing a wide variety of automated training experiments on non-neural mammalian cells, using timed drug pulses as the stimulus, and a mobile fluorescence microscope to capture images of responses, across replicate cultures. The Cell Trainer can operate in either an open-loop (feedforward) or closed-loop (feedback-controlled) mode, and our image analysis pipeline can report the behaviors of individual cells throughout each experiment and quantify population heterogeneity. We showcase the ability of the Cell Trainer to execute experimental protocols and perform single-cell analyses in both modes. We first demonstrate with a feedforward experiment in which myoblasts are repeatedly pulsed with dimethyl sulfoxide (DMSO) and their discrete calcium responses are analyzed, revealing sensitization-like dynamics. Next, we demonstrate a feedback control scheme wherein the fluorescence of a pH/voltage reporter in kidney cells is maintained below a threshold level with controlled pulses of acid. To accelerate research in the field of cell training, learning, and memory, we are openly sharing the Cell Trainer schematics and software with the research community. This platform provides a flexible tool for studying how cellular physiological states can be shaped by patterned stimulation and feedback control through approaches that work with the native adaptive competencies of cells.

## 1. Introduction

### 1.1 Background

Despite our growing knowledge of molecular biology, the task of controlling the physiology of cells remains challenging. In part, this is due to cells’ complexity, making the tasks of deciphering accurate models of regulatory networks and using them to infer effective interventions difficult. Less commonly appreciated is the challenge posed by the regulative capabilities that cells possess to work against imposed stimuli and challenges. For example, cancer cells are adept at thwarting medical interventions by adapting to resist chemotherapies [1], leading to approximately 90% of cancer-related deaths [2]. Cells adapt to diverse drugs, genetic manipulations, or environmental stressors without the need for genetic mutation and selection [3–9], being competent at solving problems on-the-fly in the transcriptional, metabolic, and physiological state spaces they traverse [10].

Further study of cell responses has shown that they possess behaviors that functionally resemble classical forms of learning [11–13]. Habituation, which is considered simple learning in which an organism reduces its response in the face of repeated stimulation without exhaustion, has been observed in the unicellular *Stentor coeruleus* [14], human embryonic kidney (HEK293) cells [15] and adrenal gland tumor (PC12) cells [16, 17], and the unicellular slime mold *Physarum polycephalum* [18]. Associative conditioning (a.k.a. classical or Pavlovian conditioning) has been reported in *Stentor coeruleus* [19], *Capsaspora owczarzaki* [20], and possibly *Paramecium caudatum* [21, 22] and several amoebae [23]. Recently, HEK293 cells were shown to exhibit the mass-spaced effect (a hallmark of memory formation), and the inhibition of factors known to be critical to memory formation in brains (CREB and ERK) blocked the effect [24]. In addition to these strictly-defined forms of learning, cells have been found to exhibit more general problem-solving and memory-retaining competencies. Yeast cells have been repeatedly shown to search the space of gene expression profiles to adapt to novel problems in an exploratory manner [6, 7, 9, 25–27], and can transiently store information in the dynamics of their gene expression and chemical reactions [28, 29], even utilizing a form of reservoir computing [30], a feat typically thought relegated to nervous systems [31]. Several studies suggest that behaviors learned by an animal may be preserved despite the loss or lack of synaptic connections, or can transferred to other animals through single-cell [32] or even molecular [33] bottlenecks, demonstrating that engrams of conventional forms of memory can exist outside of brains, in some cases in single cells [34–44]. Our lab has previously shown that simulated gene regulatory networks (GRNs) exhibit several forms of memory [45, 46], which furthermore can be analyzed with metrics used to probe neural network integration [47]. Models of real GRNs exhibited more instances of memory than random networks, and behaviors could be trained with stimulation-based techniques that do not require network rewiring. Other studies [48] of cell learning and its hypothetical underlying mechanisms [49–56] are ongoing.

More broadly, we and others have argued that these capacities offer an attractive target for therapeutics [57–62], by using training techniques from behavioral science as a top-down approach to controlling their physiology, complementary to existing bottom-up network rewiring approaches of molecular, synthetic biology [63–65]. Humans have trained animals for thousands of years without any understanding, or direct manipulation, of their underlying neurobiology; by analogy, we hypothesize that these same techniques may be ported to work with the adaptive competencies of cells to shape their behaviors through experiences, without the need to micromanage their underlying molecular networks (Figure 1A), and without a complete understanding of all the internal mechanisms and their parameters. Here, we report the first of our efforts to design and implement open-access, automated, closed-loop systems for investigating cellular competencies.

**Figure 1.**
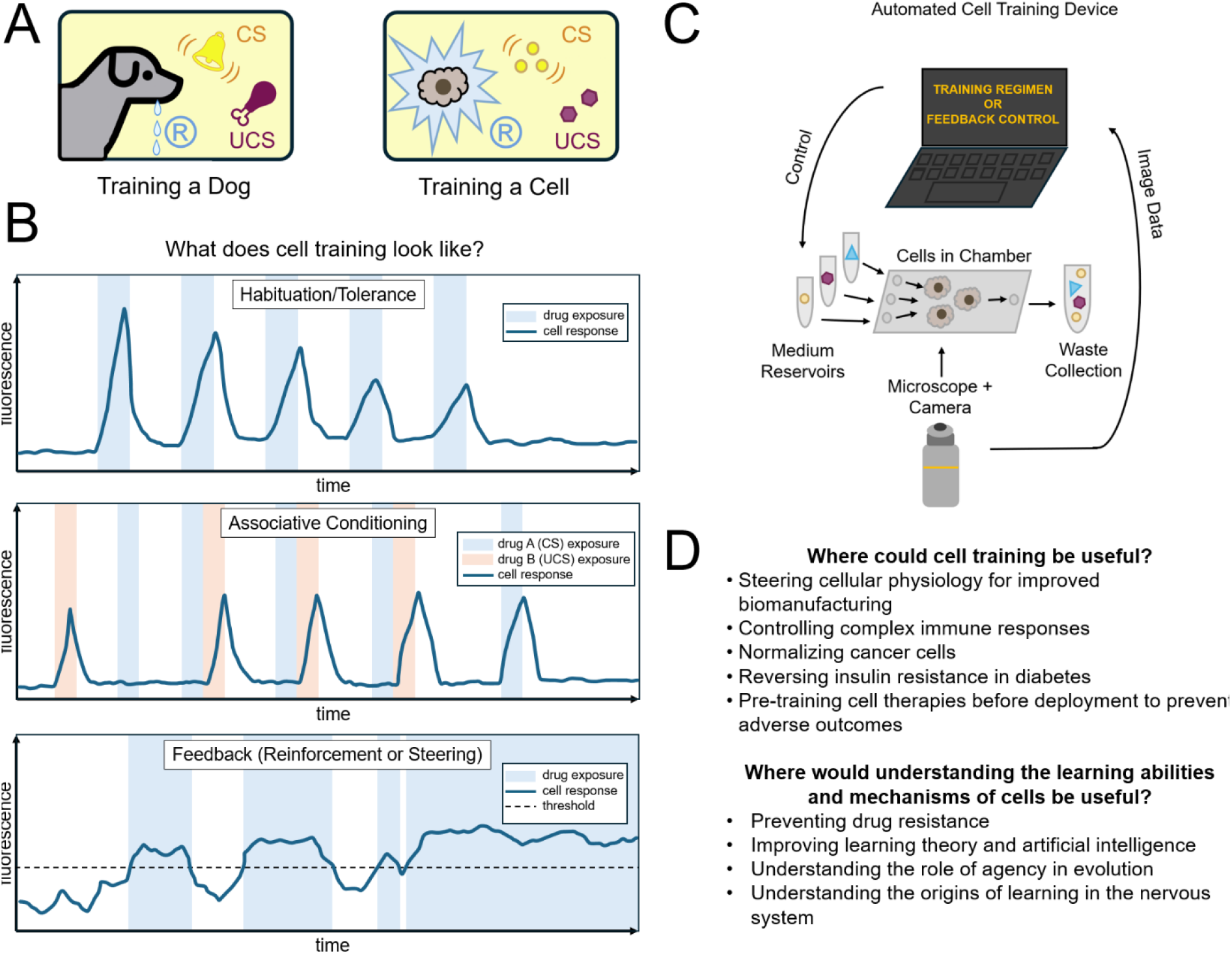
A schematic of cell training and its implementation. (A) The typical approach to controlling cell physiology is to rewire pathways. Given evidence that cells and subcellular networks can learn, we suspect that the techniques of behavioral science will be useful for controlling the physiological behaviors of cells by providing experiences that stably change behaviors without micromanaging underlying pathways, analogous to training an animal. (Image reproduced from [47].) (B) One way of conducting cell training experiments in vitro is to use drug pulses as the stimulus and fluorescent reporters of physiology as the response. (Top) A habituation experiment would consist of evenly-spaced pulses of a drug that elicits a measurable response, the magnitude of which decreases with subsequent pulses. (Middle) To train cells to associate an initially inert drug with a potent one, the inert drug may be repeatedly given before the potent drug, eventually causing the cell to use the inert drug to anticipate the potent one and respond to it as if it were the potent drug. (Bottom) A system with feedback control that can adjust the drug input based on cell behaviors could enable reinforcement learning (operant conditioning) through rewards and punishments, or model-based “steering” of cell states. (C) A device capable of performing these experiments would consist of a fluidic cell culture chamber with computer-controlled perfusion of drugs over time, while a fluorescence microscope continuously records the cell responses, which can then be used for feedback control. (D) Cell training as a novel technique for controlling cell physiology could open new doors in bioengineering and medicine.

### 1.2 A new platform

To explore the efficacy of training techniques and the limits of learning in mammalian cells, a platform for running automated training experiments is needed; previously published experimental systems are not optimized for this task. Training of the contraction behavior of *Stentor coeruleus* cells was automated with a mechanical tapping device and a camera [14, 19], but such a device is limited to a single, mechanical stimulus modality, and human cells lack this contractile response. Networks of human neurons have been trained using microelectrode arrays (MEAs) that both measure action potentials and provide electrical stimulation to cells in a closed-loop manner [66, 67], but these do not interface well with the slow bioelectric behaviors of non-excitable cells [68–70], and such techniques are designed for training networks, not single cells. The chemical switching method used to observe the mass-spaced learning effect in human HEK293 cells could be adapted for studying pulses of other kinds of drugs, but does not scale well as it requires manual delivery of drug pulses [24]. An automated training device with closed-loop control is essential to potentiate advances in both the field of diverse intelligence [71, 72] and therapeutics by enabling behavioral experiments to interface with the rise of recent lab automation [73–75] and AI tools [76] in the life sciences.

To address these limitations and opportunities for several fields, we developed a platform capable of conducting automated training experiments that closely resemble the experiments of the computational GRN training studies [45, 46], using real pulses of drugs or other chemicals as the stimuli and reporters of physiological parameters as the responses, and to perform them on mammalian cells (hypothetical cell training protocols and responses are illustrated in Figure 1B). A device capable of performing these experiments would resemble the diagram in Figure 1C, featuring cells housed in fluidic chambers with computer-controlled, timed dispensing of several drugs into the cell culture and continuous monitoring of physiological states of individual cells using fluorescent reporters and microscopy, with optional feedback control for closed-loop experiments.

In this paper, we present a device, the Cell Trainer, for automated cell training experiments using timed drug exposures as the stimulus, fluorescence microscopy as the behavioral readout, and the option to conduct either feedforward or feedback control experiments. Further, we present an image analysis pipeline capable of tracking the fluorescence levels of individual cells across the time course of an experiment, revealing heterogeneity in the learning cell population. We showcase example results from feedforward and feedback control experiments demonstrating the ability of the Cell Trainer to deliver a pre-scheduled or adaptive set of drug pulses, respectively, and to measure and perform post-hoc analysis on the single-cell responses over the course of the experiment. We are publicly disseminating the hardware design and software for the Cell Trainer to allow other scientists to make use of the device and accelerate the cell learning field and its applications (Figure 1D).

## 2. Results

### 2.1 The Cell Trainer hardware

The Cell Trainer consists of a microfluidic perfusion plate for timed delivery of drug pulses to cell cultures, a mobile microscope for repeated fluorescence imaging of multiple cultures within the plate, and a computer for control (Figure 2). The microfluidic perfusion system is the CellASIC® ONIX2 (Figure 2A, B). On each plate, cells can be housed in 4 parallel chambers each supplied by 4 medium reservoir wells. The CellASIC® ONIX2 controller enables medium switching in the chambers via air pressure-driven flow. Depending on the flow rate, the CellASIC® ONIX2 can perfuse cultures continuously for several days and can easily be refilled with medium to continue experiments longer. The mobile microscope is a custom optical system mounted on motorized translation stages (Figure 2A, C). During experiments, it moves between predefined imaging positions in a cycle, and can take fluorescence images with blue (470 nm), green (565 nm), or red (625 nm) excitation, and white-light brightfield images, at each location. A machined platform holds the CellASIC® ONIX2 plate above the microscope, and has adaptors to allow the use of other standard-sized microfluidic chips. By employing a moving microscope and static platform, this design avoids unwanted disturbance of the culture medium and fluidic components that could have unintentional impacts on cells’ exposure to chemicals and shear stresses. Schematics for the optical system can be found in the Supplemental Materials. The CellASIC® ONIX2 plate and the microscope are housed fully within an incubator that maintains a 5% CO_2_ atmosphere at 37 °C. The incubator is not humidified in order to prevent damaging the electronics, and the CellASIC® ONIX2 plate’s vacuum-sealed lid prevents medium evaporation. Software that controls the microfluidics and the microscope (described below) run on a laptop outside the incubator.

**Figure 2.**
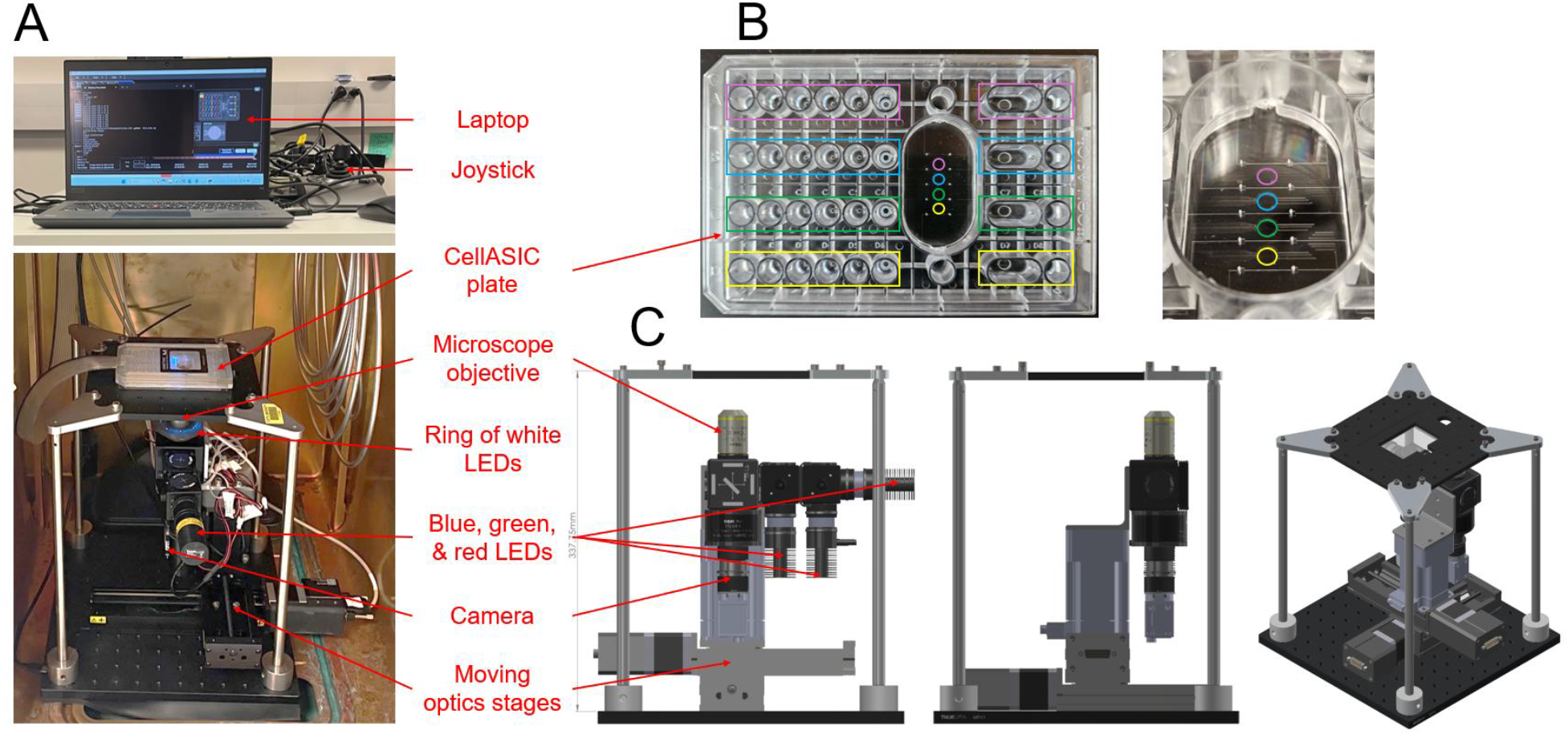
The Cell Trainer hardware can automatically deliver timed drug pulses and measure responses with fluorescence microscopy. (A) The device is primarily housed in an incubator and consists of a CellASIC® ONIX2 microfluidic plate for perfusion cell culture and a moving microscope for repeated imaging of multiple locations in the plate. A laptop and CellASIC® ONIX2 controller outside the incubator control the fluidics and optics. (B) Each CellASIC® ONIX2 plate (left) is the size of a standard 96-well plate and has four microfluidic cell culture chambers (right) that can be perfused simultaneously, and has four medium reservoirs per chamber that hold different drugs. (C) A custom-built moving microscope can repeatedly take images of multiple locations in the CellASIC® ONIX2 plate in three fluorescence channels (blue (470 nm), green (565 nm), and red (625 nm) excitation), or with white light illumination, using a Nikon 10X Plan Fluorite imaging objective and a2A4504-um18 Basler camera. A joystick next to the laptop is used to position the microscope.

### 2.2 The Cell Trainer software

The microscope of the Cell Trainer is controlled by custom software (Figure 3A). For each experiment, the microscope moves in a loop between pre-defined imaging locations to produce an image time series for each location. Once the cell cultures are in place, prior to starting the experiment, the microscope is manually moved into the desired imaging positions with a joystick, and their x, y, and z coordinates are saved, and the imaging time interval is defined. Optimal imaging settings are determined manually and saved (camera exposure and gain, illumination intensity). The CellASIC® ONIX2 software is used to specify the stimulus schedule (Figure 3B). Within a CellASIC® ONIX2 experiment file, the user specifies a fixed series of time intervals during which a specified set of medium reservoirs (i.e., drugs or chemicals) are perfused through the cell chambers at specified dispensing pressures. For “feedforward” (open-loop) experiments in which the cells are to be exposed to a pre-defined schedule of drug pulses, a single CellASIC® ONIX2 experiment file is executed while, simultaneously, the microscope imaging loop runs and continuously collects images (Figure 3C). For “feedback” (closed-loop) experiments, where the device must adapt the drug delivery schedule based on the behaviors of the cells, a set of CellASIC® ONIX2 experiment files representing each perfusion state of interest are defined prior to the experiment. During the experiment, an additional controller runs continuously, processing the latest image taken by the microscope and determining which media should be dispensed. This decision is sent to the CellASIC® ONIX2 software, closing the current experiment file and running the one representing the appropriate state.

**Figure 3.**
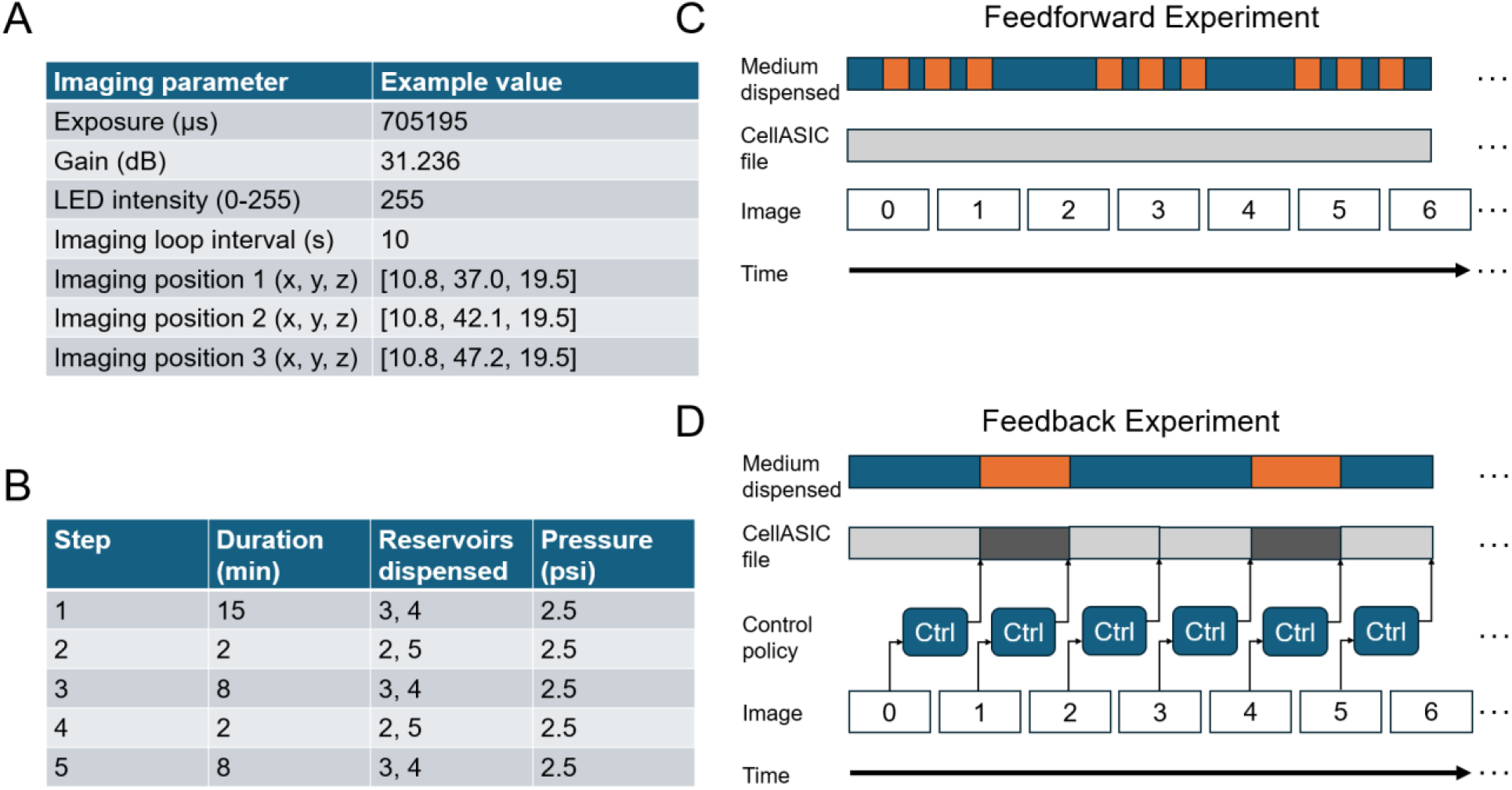
The Cell Trainer software can be configured for a variety of experiments. (A) An imaging loop is defined for each experiment, including imaging position coordinates for multiple locations, imaging interval, and lighting and camera settings. (B) The fluidics are controlled by the CellASIC® ONIX2 software which delivers medium from specified reservoirs at pre-specified times. (C) Feedforward training experiments can be performed by running a single CellASIC® ONIX2 file with pre-defined solution switching times with a simultaneous imaging loop. (D) For feedback control experiments, an additional controller process runs that processes each latest image to determine cell behaviors and, based on the control policy, determines which drug should be dispensed by sending commands to switch CellASIC® ONIX2 files. The closed loop time is less than 1 second.

A custom pipeline was written for post-hoc analysis of images from the Cell Trainer (Figure 4). For each imaging location (i.e., each cell culture chamber), a series of images is generated from the experiment. First, the cells in each image are identified in one of two ways: If the cells do not move during the experiment, one representative image is chosen and individual cells are identified via semi-manual segmentation using the Cellpose software [77]. This cell mask can then be applied to all images in the dataset. Alternatively, if cells do move during the experiment, segmentation parameters are first found for one representative image and then applied to produce a mask for each image in the series. Then, cell identities are tracked across the series based on the amount they overlap from one image to the next. Brightness artifacts caused by optical imperfections that are uneven across space and fluorescent medium components that change across time are present in the images and may impact apparent cell fluorescence levels. For each image, the background artifact is estimated and subtracted. The fluorescence of each cell is tracked across the image series and then normalized using ΔF/F0 normalization. Once the normalized single cell traces are produced, they can be analyzed further, e.g., to quantify learning and its heterogeneity across the population, to look for spatial relations between cell behaviors, to identify behavioral subtypes, and other analyses of interest.

**Figure 4.**
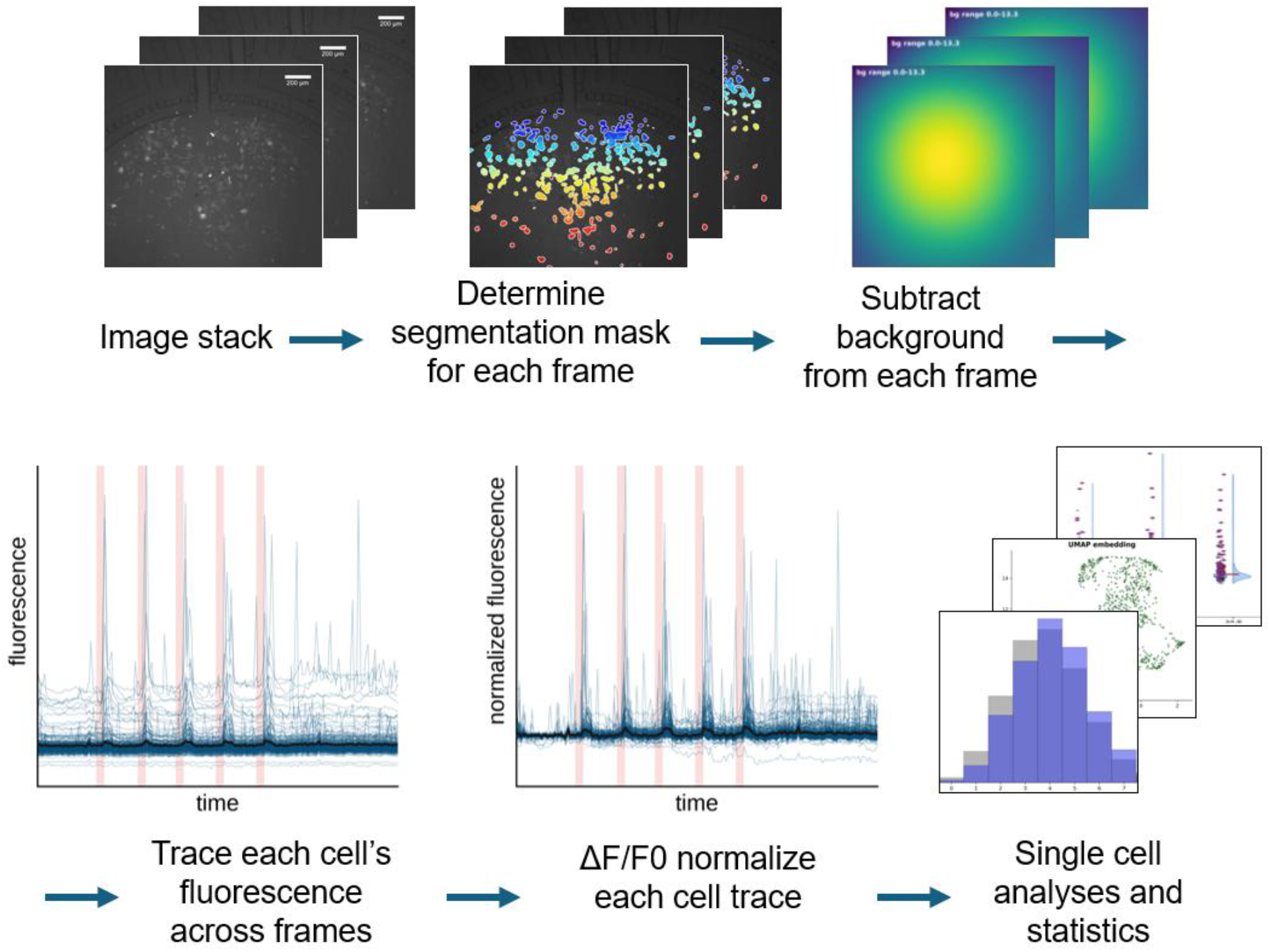
The post-hoc image processing pipeline of the Cell Trainer enables tracking and analysis of individual cell behaviors. A custom pipeline was written for post-hoc single-cell analysis after the completion of an experiment. A cell segmentation mask is determined for each frame using a semi-manual process in which the user finds the optimal segmentation parameters in the Cellpose software, and the background brightness artifact is estimated and subtracted. If cells move significantly during the experiment, their positions can be tracked automatically across frames. The fluorescence intensity of each cell is traced across the experiment and normalized. The cell population can then be analyzed for heterogeneity (e.g., good vs. poor learners), spatial patterns, etc.

### 2.3 Feedforward experiment: repeated DMSO induces discrete GCaMP6f responses

We tested the ability of the Cell Trainer and its analysis pipeline to perform a feedforward experiment and to quantify single-cell behaviors (Figures 5-7). The experiment uses C2C12 cells (a mouse skeletal muscle myoblast cell line) expressing the genetically encoded cytoplasmic calcium reporter, GCaMP6f, which increases its fluorescence with increasing cytoplasmic calcium levels (Figure 5A). We found that, upon exposure to a pulse of medium with 5% dimethyl sulfoxide (DMSO), GCaMP6f flashes brightly [78, 79]. If the DMSO is removed and another pulse is given, the cells flash again. We prepared a CellASIC® ONIX2 plate with C2C12 cells in 3 chambers, along with normal perfusion medium, and medium containing 5% DMSO. While being repeatedly imaged every 21 seconds, cells were first perfused with perfusion medium for 10 minutes, followed by 2 minutes of DMSO medium and 8 minutes of perfusion medium looped 5 times (constituting 1 stimulus train). Cells were then given 30 minutes of perfusion medium to rest followed by another train of 5 DMSO pulses, another 30-minute rest, and a final train of 5 DMSO pulses.

**Figure 5.**
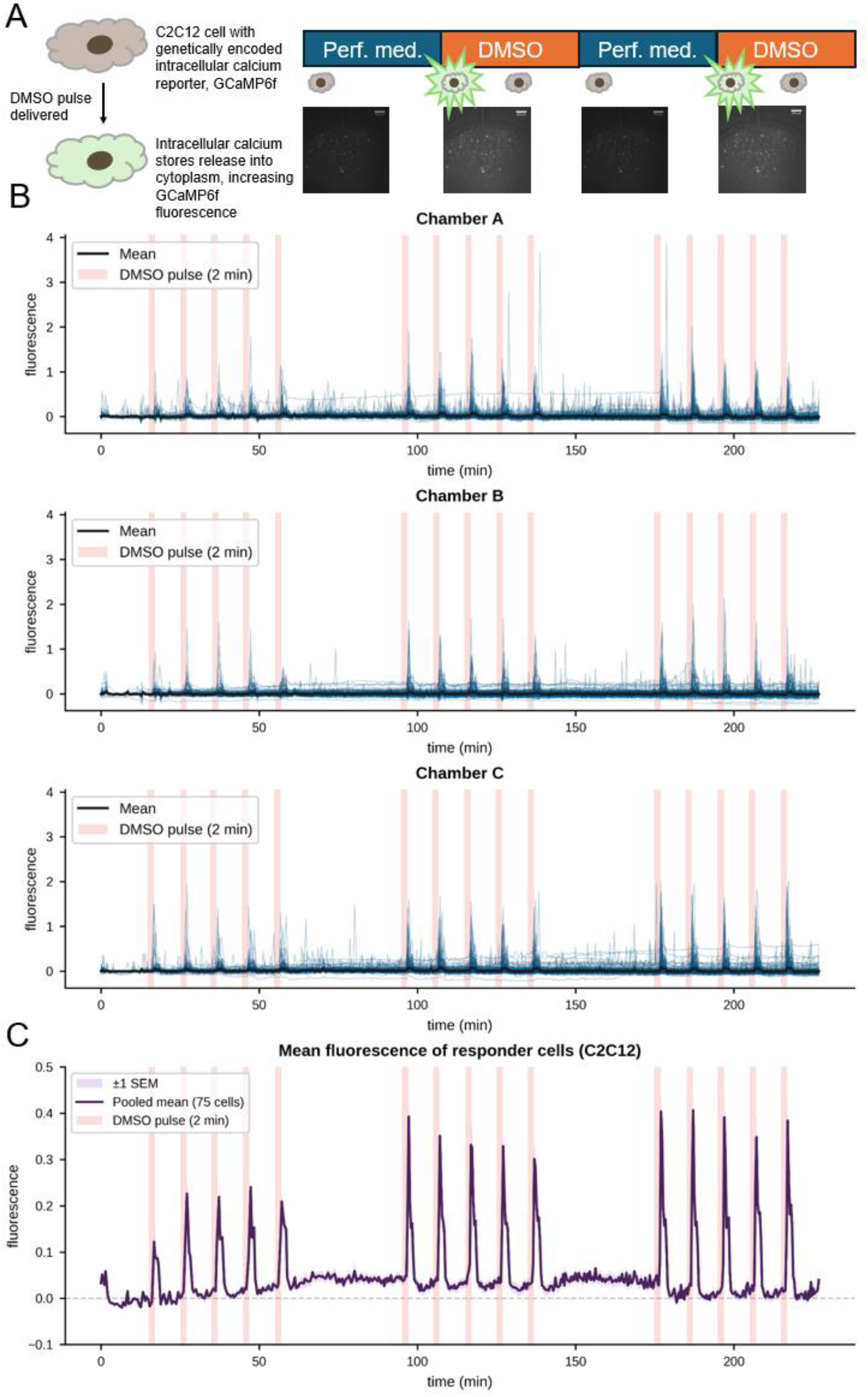
The Cell Trainer successfully performs feedforward experiments and tracks single-cell behaviors. (A) It was found that exposure to DMSO causes a minutes-long spike in cytoplasmic calcium in C2C12 cells that is visualized by the genetically encoded calcium reporter, GCaMP6f. Switching between normal perfusion medium and 5% DMSO medium induces repeating spikes. (B) In three replicate CellASIC® ONIX2 chambers, cells were exposed to three stimulus trains (i.e., series) of five two-minute pulses of DMSO every ten minutes, with thirty minutes in between stimulus trains. Each plot represents one chamber and shows the normalized fluorescence trace of each individual cell (blue lines) overlaid with the average trace (black line) and the times during which DMSO medium was delivered (red shading). (C) “Non-responder” cells were filtered out from the three chambers the responders were pooled, and their average plotted to reveal the typical cell response to the stimuli.

Figure 5B shows the normalized fluorescence traces of the individual cells in each culture chamber, with the average trace overlaid in black, and the DMSO dispensing periods shaded in red. “Non-responder” cells (cells lacking response magnitudes significantly different from baseline fluctuations; see Materials and Methods) were filtered out, and the remaining cell traces were pooled, averaged, and plotted in Figure 5C. Cells exhibited a clear calcium response to each stimulus. Interestingly, each peak appears to have a “notched” shape, explored more in Figure 8, and appears to undershoot before increasing to its final steady value following the stimulus. It also appears that response magnitudes tend to increase with each subsequent stimulus train, which is analyzed in Figure 6 C and D. Thus, Figure 5 demonstrates the ability of the Cell Trainer to run a feedforward experiment consisting of chemical pulses eliciting discrete fluorescence responses from cells (suitable for, e.g., habituation and associative learning experiments), and for the analysis software to resolve and distinguish the behaviors of all individual cells in the chambers, enabling investigations into variability in behaviors and learning abilities across cell populations. Movies of the C2C12 chambers can be found in the Supplementary Materials, and segmentation masks and raw and normalized fluorescence traces of each cell can be found on the project GitHub repository.

**Figure 6.**
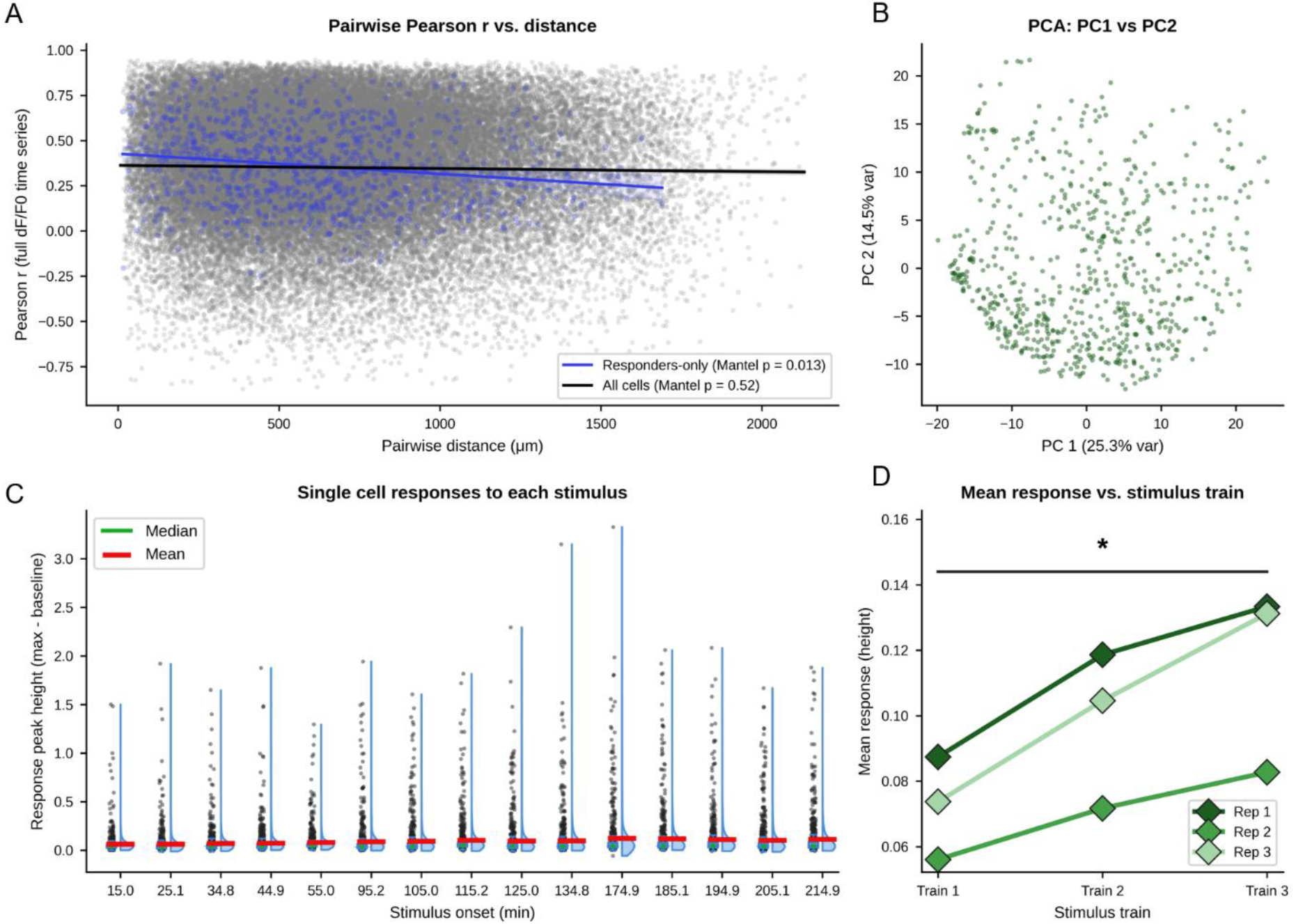
Analyses can be performed on single-cell behaviors. (A) Potential spatial relations between cells’ behaviors can be uncovered. For each pair of cells in a given chamber, the Pearson correlation coefficient, r, was found, along with the distance between the pair of cells. For all cell pairs pooled (gray and blue dots, black line), from 3 chambers (biological replicates) the line fit for the scatterplot showed a weak correlation-distance relationship (slope = −1.74e-05 Δr/μm, fit r = −0.021, 81766 cell pairs; replicate-level Mantel p = 0.52 (n = 3 cell culture chambers, mean per-chamber r = −0.019)), while the results for pooled responder cells alone (blue dots, blue line) showed a slightly stronger and statistically significant negative relation between r and pairwise distance (slope = −1.12e-04 Δr/μm, fit r = −0.168, 1047 cell pairs; replicate-level Mantel p = 0.013 (n = 3 channels, mean per-channel r = −0.140)). Shaded band around each fit line = ±3 SEM. (B) Dimensionality reduction techniques such as principal component analysis (PCA) can be used to plot cells to look for clusters of behavior. (C) Metrics of cell responses can reveal behavioral heterogeneity. The violin plots show response peak heights for all cells in response to all stimuli, revealing (D) a statistically significant increase in average peak height across the three pulse trains (replicate-level one sample t-test between first and third train, p = 0.0404).

We then performed several analyses to demonstrate the capacity to explore the population of single-cell traces (Figures 6 and 7). We hypothesized that cells closer to each other in space may have more similar behaviors to each other than to cells further away, possibly due to cell-cell interactions, or due to physical influences of various positions within the chambers such as fluid flow or illumination gradients. For each pair of cells in a given chamber, we computed the Pearson correlation coefficient between their traces, and the distance between the cells. The scatterplot of these values (Figure 6A) suggests that, after filtering out non-responders, there is a statistically significant negative relationship between the pairwise distance between cells and their correlation. Dimensionality reduction techniques can be applied to the cell traces to reveal clusters of behavioral types. Principal component analysis (PCA) and plotting the first two principal components of each cell, did not reveal clearly separated behavioral subpopulations (Figure 6B).

**Figure 7.**
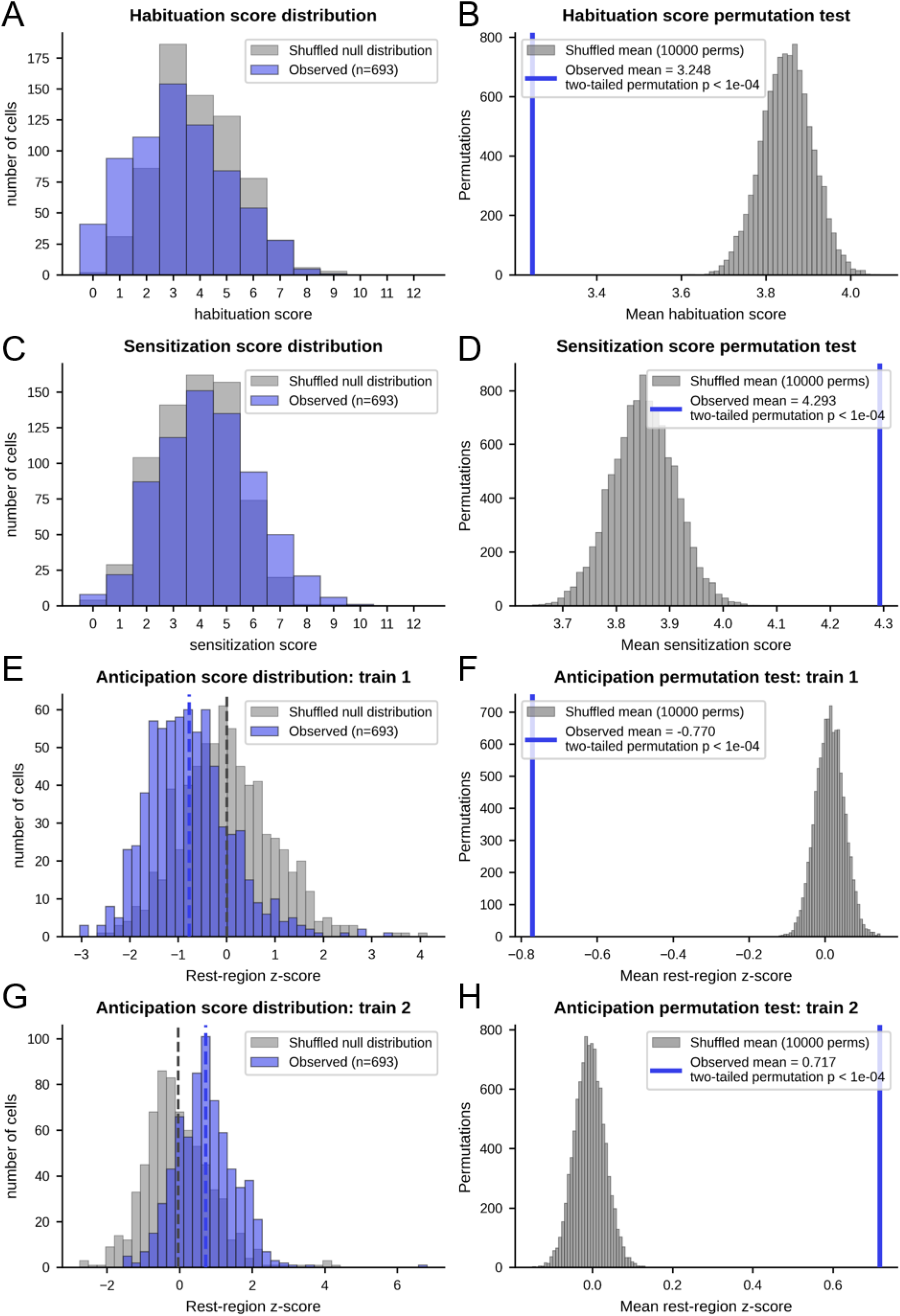
Metrics of single-cell response patterns consistent with several types of learning reveal statistically significant behavioral changes in the population. Metrics to quantify cell response patterns that are signatures consistent with three forms of learning—habituation, sensitization, and anticipation—were defined and applied to single cells (see Materials and Methods). (A) The distribution of “habituation scores” for single cell response trains is shifted towards lower values than scores for shuffled data (blue histogram shows the distribution of true single-cell habituation scores; gray histogram shows scores for single-cell data with response peak orders shuffled). (B) A permutation test reveals this difference is statistically significant. (C) A similar but converse “sensitization score” applied to the single cell responses reveals a shift towards higher scores relative to shuffled data that is statistically significant (D). The tendency of cells to anticipate a sixth pulse that never arrives can be quantified by looking for unusually high or low fluorescence values at the time when the sixth pulse would have come for each train. (E) Following the first stimulus train, cells had lower fluorescence values at the anticipation time than randomly selected values from the resting period, and the difference was statistically significant (F). (G) Following the second train, cells instead had higher fluorescence values than typical resting period values at anticipation time than typical resting period values, with a statistically significant difference (H). While these metrics alone are insufficient to declare the presence of learning, they demonstrate that the types of metrics for proper assessments of learning can be applied to single-cell responses and they reveal response trends in this experimental dataset.

In addition to analyzing cells based on their full time series traces, key behavioral metrics can be extracted from those traces and analyzed. For each stimulus, we measured the response magnitude (peak height) as the maximum value of the peak minus the value before the response, and plotted the response distribution for each peak as violin plots (Figure 6C). Tracking the mean peak height of each stimulus train (each group of 5 peaks; 3 groups total) in each culture chamber revealed a statistically significant increasing trend in average peak heights across subsequent stimulus trains (Figure 6D). Importantly, this represents a change in cellular response dynamics over repeated stimulation. Overall, Figure 6 shows that the measured population of single-cell behaviors can be explored for patterns across space, in terms of unbiased behavioral comparisons, and in terms of behavioral metrics devised for features specific to the given experiment.

We then demonstrated that metrics of behaviors aligned with several types of learning (see Materials and Methods) can be applied to the single-cell traces from the Cell Trainer (Figure 7). Cells tended to have lower habituation scores and higher sensitization scores than the null distributions calculated from shuffled data (Figure 7 A, C, respectively). Permutation testing (Figure 7 B, D, respectively) showed that these distribution differences are statistically significant, suggesting that the cells tend to have larger and larger responses to subsequent stimuli within a train, and that this tendency is lost when the order of the response peaks is shuffled. Interestingly, the average fluorescence during the resting period after the first train is about equal to that of the resting period after the second train, despite the clearly rising response peaks (Figure 5C), suggesting that the increasing peaks is not simply the result of accumulating damage due to blue light exposure [78] or an apparent fluorescence increase caused by cell detachment and rounding. We then assessed whether cells exhibited unusually high or low fluorescence values at the time when they would have anticipated a non-existent sixth stimulus to arrive after a train of five stimuli (10 minutes after the last stimulus was delivered). For each of the first two stimulus trains (as the third did not have a long enough window of time following it to fit the analysis), we plotted a histogram of the normalized fluorescence values of all cells at the time they would have expected a sixth pulse to arrive (the “anticipation time”) and compared it to a histogram of fluorescence values randomly selected from the resting period of the cells (the time between stimulus trains). Interestingly, after the first train, cells tended to have lower-than-typical fluorescence levels at anticipation time (Figure 7E), while after the second train, they had higher-than-typical fluorescence levels at anticipation time (Figure 7G). Each finding was statistically significant according to a permutation test comparing the response at the anticipation time to responses at shuffled timepoints in the resting period (Figure 7 F, H). The single-cell results of the habituation and sensitization scores align with the bulk-cell result of Figure 6D, showing an overall tendency for behavior change during the experiment, while also making clear the heterogeneity amongst the cell population. In summary, we show that learning type-specific metrics can be applied to single-cell responses Furthermore, we showed that the GCaMP6f responses of C2C12 cells to DMSO pulses have a statistically significant tendency to increase with subsequent pulses, which is consistent with sensitization, while also revealing apparently opposing tendencies to anticipate non-existent stimuli, requiring further study.

The Cell Trainer can resolve subtle behavioral differences between different cell lines under replicate experimental conditions. The C2C12 experiment above was repeated with 1 cell culture chamber containing a different cell line (Figure 8A). PC-3 cells (a human prostate cancer line) expressing GCaMP6f exposed to the same DMSO pulse schedule as the C2C12 experiment had several differences compared to the C2C12 cells. First, their signal was less noisy due to a lower amount of spontaneous calcium activity in the cells in between stimuli. Second, the shape of their response peaks was more rounded, lacking the “notch” seen in the C2C12 peaks (a biphasic response), suggesting different underlying mechanisms driving calcium release and sequestration dynamics, warranting further study. A movie of the PC-3 chamber can be found in the Supplementary Materials, and the segmentation mask and raw and normalized fluorescence traces of each cell can be found on the project GitHub repository.

**Figure 8.**
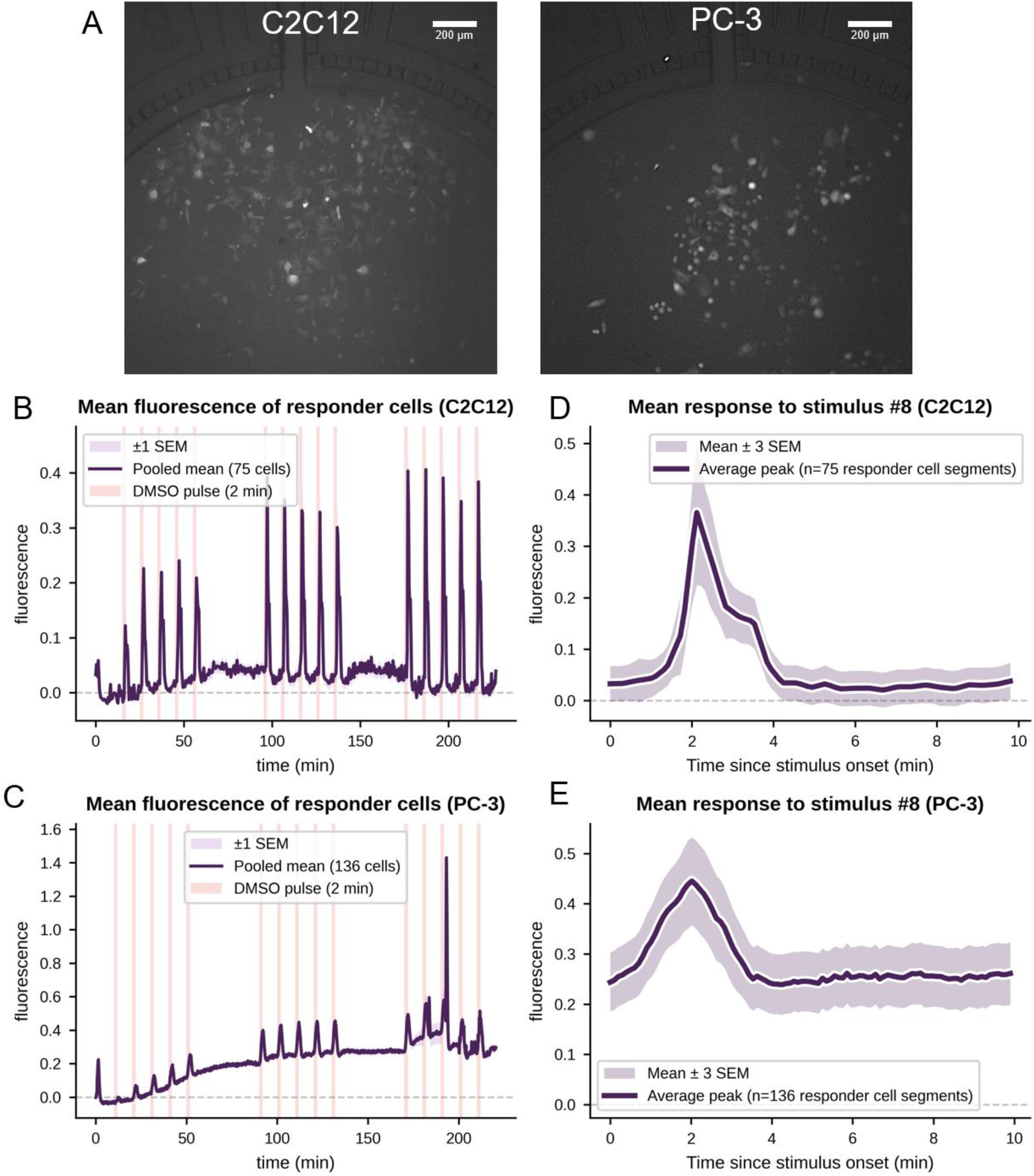
The Cell Trainer reveals cell type-dependent differences in behaviors. (A) C2C12 cells and PC-3 cells were subjected to the same DMSO stimulation schedule in separate experiments in the CellASIC® ONIX2 (scale bar = 200 μm). (B) C2C12 cells exhibited more GCaMP6f “flickering” throughout the experiment, resulting in a noisier average fluorescence trace, and featured sharp average response peaks with a “notched” shape. (C) PC-3 cells were less noisy and had simple, round average response peaks. Subset of cells spiked extra brightly on the second and third peak of the last stimulus train, possibly reflecting cell damage, rupture, or another extreme physiological response. (D) A zoomed-in view of the third peak of the second pulse train for the C2C12 and (E) PC-3 cells, highlighting the difference in calcium response dynamics between the two cell types.

### 2.4 Feedback experiment: controlling ArcLight with pulses of acidic medium

We demonstrated the ability of the Cell Trainer to perform feedback control experiments, independently controlling two chambers simultaneously. We used NRK-49F cells (a rat kidney fibroblast cell line) expressing the genetically encoded pH/voltage indicator, ArcLight (Figure 9A). In normal perfusion medium, the cells fluoresce constantly, but upon exposure to acidic medium, the fluorescence decreases rapidly [80], then slowly increases upon returning to perfusion medium. The objective of the controller in this experiment was to keep the average cell fluorescence below a setpoint defined as approximately 1 fluorescence unit below the initial value during experiment setup. The control policy was to deliver normal perfusion medium by default, but if the device detected that the average fluorescence exceeded the setpoint, it delivered a 30 second pulse of acidic medium, causing a decrease in fluorescence intensity. (Figure 9B) The experiment began with 5 minutes of normal perfusion medium and the controller turned off. When the controller was turned on, the device began to compute the average cell fluorescence in each image shortly after it was taken, made a decision about which medium to dispense, and sent its commands to the CellASIC® ONIX2 fluidics.

**Figure 9.**
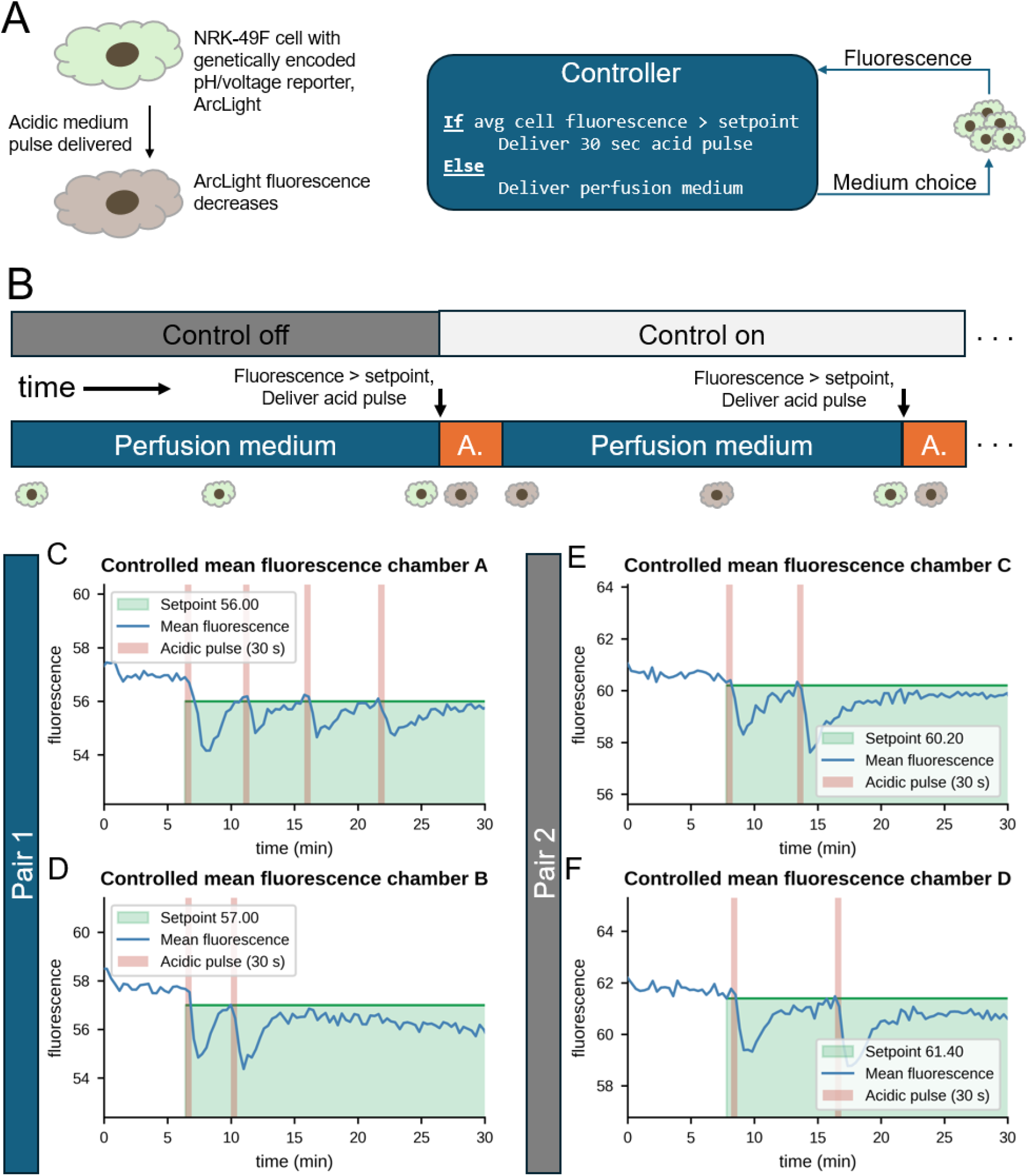
The Cell Trainer successfully performs feedback experiments to control cell physiology in pairs of cell culture chambers simultaneously. (A) NRK-49F cells expressing the genetically encoded pH/voltage reporter, ArcLight, are continuously fluorescent in normal perfusion medium, but rapidly decrease in fluorescence upon exposure to acidic medium. A control policy was implemented such that, if the average cell fluorescence in a culture chamber is greater than a setpoint, the device delivers a 30 second pulse of acidic medium, decreasing the fluorescence. (B) Experiments began with control off for 5 minutes, allowing cells to acclimate, after which control was turned on. A pair of culture chambers could be controlled simultaneously, and Pair 1 (C, D) and Pair 2 (E, F) were tested in separate experiments. Each plot shows the average cell fluorescence of a given chamber (n = 1 chamber) over 30 minutes, with control beginning after 5 minutes, with the setpoint shown by a green horizontal line. Red shaded areas indicate the times during which acid pulses were being delivered. In each chamber, the controller successfully sends a pulse each time the average cell brightness surpasses the setpoint. Note that after multiple exposures, several cells died due to acid exposure, lowering the average fluorescence to the point that it stopped exceeding the setpoint.

Figure 9 C-F shows the average cell fluorescence traces for two pairs of two simultaneously controlled chambers. In each, once control is turned on after 5 minutes, the device successfully delivers an acid pulse (red shaded area) each time the average fluorescence exceeds the setpoint (horizontal green line), causing the fluorescence to decrease before slowly returning towards the setpoint. Note that, due to some cell death from acid exposure during the experiment, the population as a whole could sometimes not return to the setpoint after several acid pulses. In summary, Figure 9 proves that the device can respond to cell states in real time by updating which medium is being perfused less than 1 second after an image is taken in order to keep physiological reporter levels within a controlled range. Movies of the NRK-49F chambers can be found in the Supplementary Materials, and segmentation masks and raw and normalized fluorescence traces of each cell can be found on the project GitHub repository.

Individual cell behaviors could be resolved in feedback experiments just as they were in feedforward experiments and analyzed post-hoc. Figure 10 shows the normalized fluorescence traces of all cells (left column), which also visualizes the sub-population of cells that died and stopped responding during the experiment. Figure 10 (right column) also shows that, in each chamber, cells further apart from one another in the chamber tended to have less correlated behaviors in terms of the Pearson correlation coefficient, and that the correlation-distance relationship was statistically significant. Together, Figures 9 and 10 show that the fluidic system can respond to cell states in real time, and the behaviors of single cells can be analyzed in this context.

**Figure 10.**
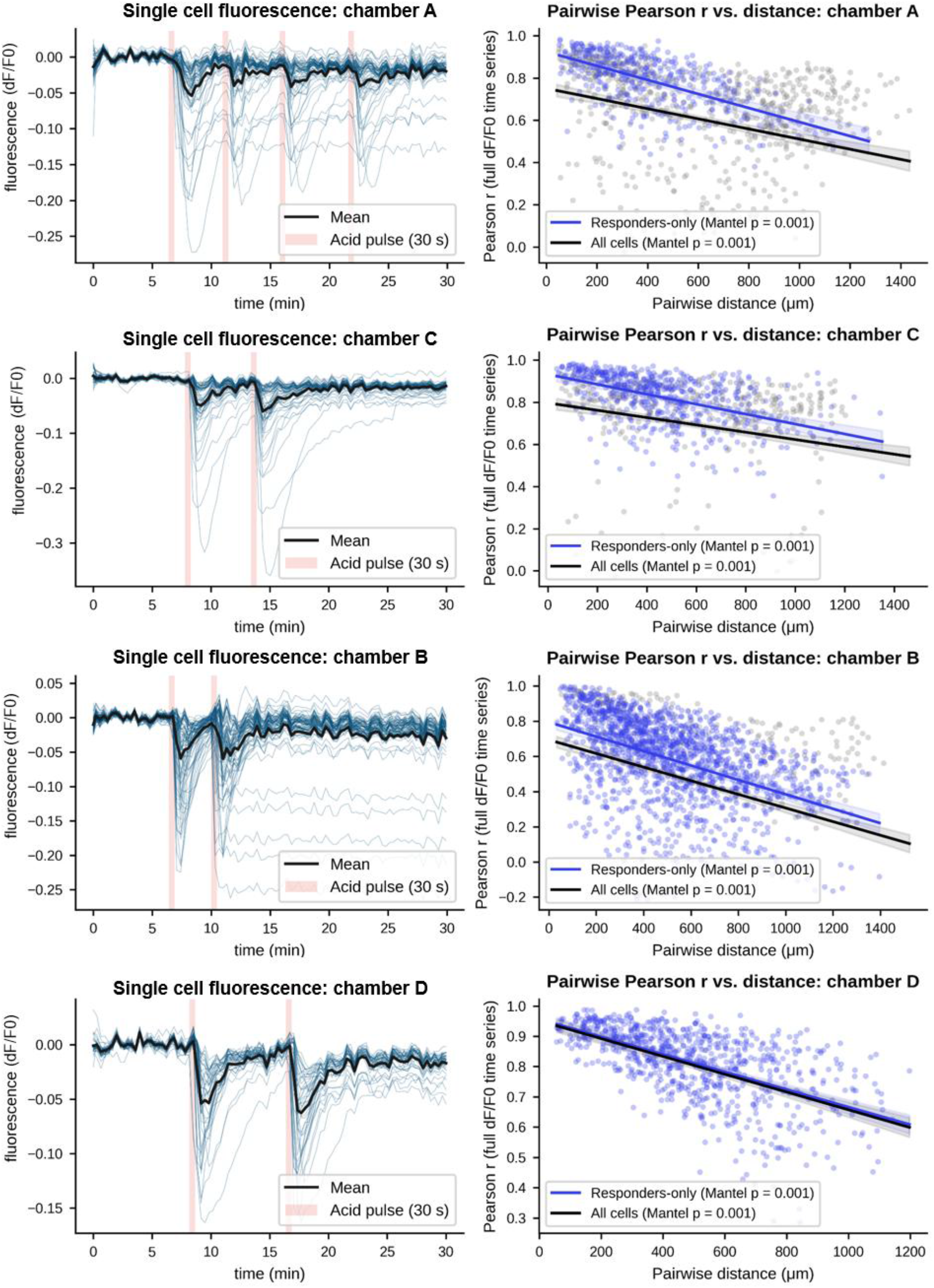
Single-cell behaviors from feedback control experiments can be visualized and analyzed. Rows show data for four different controlled cell chambers; chambers A and B were controlled simultaneously, and chambers C and D were controlled simultaneously. (Left column) The normalized single-cell traces illuminate the behaviors of all the cells in the cultures during population-level control. (Right column) Spatial relations between cell behaviors are visualized as a scatter plot of the Pearson correlation coefficient between each pair of cells in a culture vs. the distance between them, revealing that cells further apart have less correlated behaviors. Statistically significant relations (p < 0.05) were observed in all chambers when assessed for either all cells (gray and blue dots, black line), or responders only (blue dots, blue line).

## 3. Discussion

### 3.1 Summary of findings

We described new hardware, control software, and analysis software for a platform called the Cell Trainer capable of conducting automated training experiments on non-neural mammalian cells, using pulses of drugs or other chemicals as the stimuli and fluorescence microscopy to measure physiological responses of single cells. We demonstrated a feedforward experiment in which both C2C12 and PC-3 cells were stimulated with repeated pulses of DMSO and exhibited discrete calcium responses measured with GCaMP6f, the dynamics of which differed between the cell lines. We showed an analysis pipeline that can report the behaviors of the individual cells and can subject them to analyses of heterogeneity, correlation, and quantification of behaviors consistent with various forms of learning. Finally, we presented a feedback control experiment in which the device delivered pulses of acidic medium to NRK-49F cells based on real-time measurements of the mean cell fluorescence level (of the pH/voltage reporter, ArcLight) in order to keep its level below a user-specified threshold, and performed similar single-cell behavioral analyses post-hoc.

While our biological findings are preliminary (full analyses of cells’ learning capacity will be presented in future publications), the increasing response of GCaMP6f to repeated DMSO pulses is consistent with sensitization, a form of learning in which the magnitude of a response increases with repeated stimuli. This occurred without an apparent rise in baseline GCaMP6f fluorescence, suggesting the increasing response is not simply due to cumulative damage or cell morphology change. The differences in the shapes of the response dynamics across the two cell lines tested with DMSO point towards differences in the underlying mechanisms each uses to release and sequester its internal calcium stores, an unexpected finding that should be investigated to further our understanding of calcium biology. In addition to proving that the device can perform closed-loop control, the experimental manipulation of the ArcLight reporter with pulses of acidic medium also revealed an apparent asymmetry in ion fluxes across the NRK-49F cell membranes. Specifically, upon the sudden switch from normal to acidic medium, the cells’ fluorescence decreases rapidly. In contrast, when medium is suddenly switched from acidic to normal, ArcLight takes much longer to return to its initial value. This may indicate that the configuration of the cells’ ion pumps and channels greatly favor influx of protons over efflux, although future experiments are needed.

### 3.2 Future improvements to the platform

The current hardware of this v1.0 Cell Trainer platform will be improved in future work to expand its capabilities. The optical system can be augmented with the addition of phase contrast microscopy, and by improvements in the microscope movement speed, which currently results in a ∼20 second minimum time interval between images if three CellASIC® ONIX2 chambers are used. Alternative fluidic systems may be used to provide additional chambers and reservoirs, independent control of reservoirs across chambers, or chambers of alternative designs that can accommodate different biological entities (e.g., organoids) or flow patterns. The current system can produce step changes in chemical concentrations; instead, a continuous range of chemical concentrations might be produced using pulse width or pulse frequency modulation [81], and pharmacokinetic profiles may be modeled with the incorporation of mixing chambers [82]. New sensors and stimulus modalities could be added to expand the range of inputs and readouts. Optogenetic, electrical (such as with micro electrode arrays), thermal, and other forms of inputs and readouts could be added, and existing hardware may be repurposed, such as using the fluorescence excitation LEDs for optogenetic stimulation. Shear stress may also be explored as an input in the current system, controlled by changing flow rates. Without the addition of any new sensors, the current device may be used to measure not just fluorescence, but any other feature extractable from image data in the 4 available channels, including cell shape and movement. Alternative microscopy systems could enable rich, label-free readouts of chemical compositions with various hyperspectral or spectroscopic techniques [83–87], or measurements of the chemical composition of effluent medium from the cell culture to track their secretion or absorbance rates of substances of interest. Effluent medium may be fractionated, collected, and chemically analyzed to resolve its composition over time [88, 89]. Improvements can be made to the imaging loop software to enable automated capture of videos of cells as opposed to only still images. An improved user interface will be developed to integrate and streamline the use of the imaging software together with the feedback control software. In the current image analysis pipeline, the automated cell segmentation can run slowly on large datasets, requires manual optimization of parameters, and imperfect segmentation may inhibit proper tracking of cells. While no segmentation software can ever work perfectly, segmentation software is constantly improving and will be updated in our pipeline.

### 3.3 Future experiments and applications

The purpose of this paper was to demonstrate a device capable of performing cell training experiment protocols and an analysis pipeline for making sense of single-cell behavioral outcomes. While we did observe statistically significant response increases in C2C12 cells that are consistent with sensitization, additional defining features of sensitization (defined the same as habituation [90] but for increasing responses) were not tested or quantified. Furthermore, apparent anticipation was observed in C2C12 cells after each of the first two stimulus trains, but their values were in opposite directions, requiring additional interpretation. Finally, while many individual cells were included in the analysis, they were from only 3 replicate chambers for C2C12 experiments, 1 chamber for the PC-3 experiment. For the NRK-49F experiments, only 4 chambers (2 pairs) were used, and each chamber was analyzed separately. Future biological studies with claims of cell training and cell learning will be conducted with larger numbers of replicates to ensure robust conclusions and will assess learning against thorough definitions. Further experiments are needed to understand the mechanisms and nature of the behavioral change we observed in DMSO responses, including its persistence, its specificity to the DMSO stimulus, and to contextualize it within standardized definitions of learning if simpler explanations, such as simple accumulation of membrane damage across repeated stimuli, are ruled out.

While more examples of cell learning continue to be published, the field is still in its infancy, and many specific capabilities of cells conjectured here are yet to be solidly demonstrated, such as associative conditioning in mammalian cells. Utilizing the Cell Trainer as described in this paper, we are beginning to search for forms of learning and are testing the efficacy of simple training techniques in mammalian cells. As habituation in cells has previously been described, we aim to study more sophisticated behaviors, such as associative conditioning, which has yet to be demonstrated in mammalian cells. We are also beginning to test reinforcement learning schemes where the device delivers rewards or punishments to cells to try to stably change reporter fluorescence levels. Once strong experimental models of learning are established, we can begin testing setups in which the device uses a machine learning/artificial intelligence-based controller—itself trained on the outcomes of many automated experiments [74, 91]—to automatically train cells by steering their states [80]. The feedback control system is modifiable and can implement nearly any control policy, from simple if-statements to complex machine learning based model predictive control, and it could be used to connect multiple cell cultures or other agents together, allowing them to interact [92]. The device may even be capable of incorporating large language model (LLM)-based translation and embedding of natural language commands for two-way communication with cells or other biological entities [93–95], allowing us to not only influence cell behavior with training, but to persuade them, and interpret their behaviors, via natural language. Furthermore, with an established learning model, we can characterize its features and its mechanisms, especially looking for connections to neurobiological mechanisms [24, 68–70]. We will continue evaluating reporters of different physiological parameters to study those with the most immediate medical applications. We suspect that the training principles discovered using this device should be translatable far beyond microfluidic cell culture, and we will engineer approaches to bridge these findings for use in patients, bioreactors, agriculture, and other circumstances where control of biological systems is needed. Scaled up to industrial deployment, we can envision such systems becoming part of biopharma’s screening approaches for new reagents targeting cell adaptation algorithms.

We foresee cell training becoming a technique for controlling cell physiology that is complementary to current molecular biology methods and takes advantage of cells’ adaptive competencies to enable engineers and medical practitioners to shape cell behaviors without the need to rewire underlying molecular pathways. Just as humans may be persuaded, computers may be programmed, and language models may be prompted, we hypothesize that training can serve as an appropriate “user interface” enabling efficient control of cells. In biomanufacturing, controlled stimulation in bioreactors may enable top-down control of cells to maximize production and prevent drift without tedious genetic manipulation. In medicine, training techniques may help control immune disorders and cell therapies, could steer cells from diseased states (e.g., cancerous [91, 96] or diabetic [97]) to healthy states (“forgetting” maladaptive changes [98]), or allow the body to associate the effects of a potent, yet toxic, drug with a benign one.

Improving our understanding of the mechanisms and algorithms behind cells’ learning abilities would also advance basic science. The molecular components and functional algorithms shared between neural tissues and their ancient unicellular precursors [24, 70, 99–102], may make single-cell organisms tractable model systems for studying mechanisms present in the brain. Blocking these mechanisms by translating memory blocking drugs from neuroscience may prevent the development of drug tolerance. The fields of natural and artificial intelligence have historically advanced in tandem, borrowing insights from each other; studying the learning and computational abilities of cells that we do not yet understand may lead to new breakthroughs in computational applications [103]. Standard theory in biological development, physiology, and evolution do not account well for cells that learn during their lifetimes and stand to be improved by incorporating findings from these studies [104, 105]. Finally, this research program has implications for philosophy of mind and questions about the utility of recognizing cognition in unconventional entities [71, 72].

### 3.4 Conclusion

We believe that the development of platforms like the Cell Trainer described here, and the cell training experiments they enable, serve as a first step towards a future in which system-level phenotypes in biomedical and bioengineering contexts, which may be too complex for conventional micromanaging (bottom-up molecular rewiring) approaches, are managed by shaping the history of physiological experiences, and thus behaviors, of cells, tissues, GRNs, and organs through experiences. These systems represent a powerful integration of emerging fields that include diverse intelligence [106–111], robot science [112–114], and AI [115, 116].

## 4. Materials and Methods

### 4.1 Cell Trainer device construction and software

#### 4.1.1 General device setup

The Cell Trainer consists of a microfluidic cell culture plate capable of exposing cells to drugs at specified times, paired with mobile fluorescence microscope for repeated imaging of each cell culture in the plate. These components are housed in a non-humidified incubator with a controlled atmosphere kept at 5% CO_2_ and 37 °C for mammalian cell culture. All other components are kept outside the incubator. A laptop controls the fluidics and imaging during experiments and saves the images. During experiments, the imaging loop and fluidics schedule are initiated and run independently, with the option to run a feedback controller that processes the images as they are produced and updates the fluidics schedule accordingly.

#### 4.1.2 Fluidics hardware and software

The Cell Trainer uses the CellASIC® ONIX2 Microfluidic System (MilliporeSigma/Merck KGaA, Darmstadt, Germany, CAX2-S0000) with M04S-03 microfluidic plates for adherent mammalian cells. The plate has 4 transparent microfluidic chambers to house cells. Each chamber is supplied by an independent set of 4 reservoir wells that can be loaded with various drug-containing media for the experiment. A controller unit is kept outside the incubator and connects to the plate via tubing through the incubator port to supply pressurized air to drive medium flow. The CellASIC® ONIX2 software allows users to define and run solution switching schedules, specifying which reservoirs to dispense from at what times and at what pressures.

#### 4.1.3 Microscope optics and moving stages

The custom microscope provides three fluorescence channels to the sample, as well as brightfield illumination. Brightfield illumination is provided by a custom ring light (white LEDs embedded in a 3-D printed ring) mounted around the objective. Each fluorescence channel is produced by a Thorlabs (Newton, NJ) LED (red, 625 nm, M625L4; green, 565 nm, M565L3; blue, 470 nm, M470L5) shaped by a collimating lens, with unwanted wavelengths removed by a filter before exciting the sample. The excitation channels used in this device use one of two form factors. For ease of assembly, we used Thorlabs LEDs collimated by a condenser lens (ACL25416U-A, Thorlabs), all mounted in off-the-shelf Thorlabs lens tubes (SM1L50, Thorlabs). For flatter field illumination, we used Luxeon SZ-05-H3 (Schiphol, Netherlands) Z Quad LED mounted in a custom holder and collimated by a Fresnel lens (FRP125, Thorlabs). The individual excitation beams are combined using dichroic mirrors (T525lpxr and T585lpxr, Chroma). A multiband excitation filter removes unwanted wavelengths before a multiband dichroic mirror turns the beam towards the sample (89402 ET multiband filter set, Chroma). The excitation beam is focused onto the sample by a Nikon 10X Plan Fluorite imaging objective (N10X-PF). The emitted fluorescence is collected and collimated by the objective and then filtered by a multiband emission filter. It is then focused onto the CMOS detector (a2A4504-um18, Basler) by a 100 mm infinity corrected tube lens (TTL100-A, Thorlabs).

The optical components were mounted on a custom-machined aluminum plate, mounted with m6 screws on a Thorlabs MB8 8×8 breadboard, with LSQ-150A-T3A 150mm stages from Zaber acting as x and y axes, and VSR40A-t3A from Zaber acting as the z axis is mounted on the y axis stage with the included adapter and m6 screws. These are driven by a Zaber XMCC3 Controller connected via USB to the computer. Schematics for the optical system can be found in the Supplemental Materials.

#### 4.1.4 Microscope control and image acquisition software

We developed custom software that allows the user to specify multiple imaging locations (x, y, z coordinates) and image acquisition settings. The software then moves the microscope between these locations and takes and saves images in a loop with a specified time interval. Imaging setup was implemented using a TOML file detailing imaging parameters including camera exposure and gain, LED brightness, imaging timer interval, and imaging locations (x, y, z coordinates for multiple locations) as well as through the software itself, with options for both linearly interpolated channels as well as freely added locations. The software was designed for maximum modularity and ease of adjustment, using the Python programming language to interface between stages, camera, and lights. Pypylon (https://github.com/basler/pypylon) and Zaber (https://www.zaber.com/support/docs/api/core-python/0.9.1/) APIs control the camera and stages respectively, and a custom Arduino firmware allows control of lighting over Serial communication. The nature of the software allows for different camera and stage systems to be used if necessary, with easy replacement of just those modules. Experiments are run iteratively using multiple precision timers, lowering latency to the speed of Serial communication, at 115200 Baud rate. Experiment state is saved after each imaging timepoint finishes, allowing for restarts in case of issues.

### 4.2 Feedback control software

An optional feedback controller can be used during experiments. The control script is started and runs in parallel with the imaging loop and the fluidics software, continuously monitoring the image directory to take in and process the latest image of the cells. A control policy maps the image state to cell stimulation decisions. This policy is modular and can be easily changed to meet the needs of the experiment, implementing policies ranging in sophistication from simple switches to machine learning model-based control. Once the stimulus decision is made, the controller sends commands via the CellASIC® ONIX2 API to stop the current CellASIC® ONIX2 experiment file and start the appropriate file, changing the perfusion state in less than 1 second given a sufficiently fast control policy.

### 4.3 Cell segmentation

We used Cellpose [77] for segmenting individual cells in images to produce image masks for analysis. We found that applying a median filter to images before segmentation greatly enhanced its performance (all subsequent quantitative analyses were performed on the raw image data). Attributes such as cell diameter (the size of the cells in pixels), flow threshold (the max allowed error between predicted and recomputed flows), cell probability threshold (the threshold on Cellpose’s predicted cell-probability map on a logit-like scale of roughly −6 to +6; pixels above it are assigned to cells), and NITER dynamics (the number of iterations performed to complete the segmentation) were optimized by visual inspection to produce the best segmentation. After settings were found that segmented most of the cells accurately, manual changes were made to remove non-cell regions of interest (ROIs) and to draw ROIs for any cells that were missed.

### 4.4 Image processing pipeline and single cell analyses

#### 4.4.1 Image processing pipeline general description

We created a custom Python-based image processing pipeline. For a given time series of images of cells expressing a fluorescent reporter, the pipeline identifies the cells in the images, estimates and removes the background fluorescence artifact in each image, tracks each cell’s fluorescence across the image series, and normalizes their fluorescence traces. Downstream single-cell analyses are described in the following subsections.

#### 4.4.2 Cell mask and optional cell tracking

The pipeline allows cells to be identified across the image series in one of two ways. If cells do not move during the experiment, a single, representative frame from the image series may be chosen and segmented with Cellpose. This mask is then applied to every image in the series. Alternatively, if cells move during the experiment, Cellpose settings are first tuned on a representative frame and then applied to segment every frame in the series independently. Cell identities are linked if their Euclidean separation is less than a per-experiment linking threshold (default 40 pixels; the NRK-49F acid-feedback experiment used 10 pixels). A short dropout is tolerated by a 3-frame grace period: a detection that reappears within three frames is re-linked to the same trajectory rather than being treated as a new cell. Missing frames are not interpolated; all downstream analyses use only cells detected in every frame (filter_complete_cells), so trajectories containing a gap are excluded.

#### 4.4.3 Background estimation and subtraction

Fluorescent components of the cell culture medium and imperfections in the optics introduce fluorescence patterns in the image that are uneven across space and change across time (e.g., as the medium reservoir photobleaches). To account for this, we estimate this background artifact using a polynomial fit. Sample-point locations are chosen in each imaging channel once per experiment unioned across channels of segmentation masks from 15 evenly spaced timepoints, marking every pixel occupied by a cell at any of those frames. A 25 × 25 grid of candidate sample points is placed on the image, inset by a 30-pixel margin from the edge of the frames. Each point is then displaced within a 180-pixel search radius to the location farthest from any cell, and discarded if its final clearance from the nearest cell is below 40 pixels. For each frame, the average fluorescence inside a 30-pixel radius around each point was computed, and a least-squares system for the 15 coefficients of a degree-4 polynomial was solved in normalized coordinates (x - W/2)/(W/2), (y - H/2)/(H/2). The polynomial was rendered as a full-resolution map, shifted so its minimum, evaluated on a coarse 160×160 grid, is zero, and subtracted from each cell’s measured fluorescence by evaluating the fitted surface at the cell’s tracked (x, y) at every frame.

#### 4.4.4 Producing and normalizing single cell fluorescence traces

For each frame, once the background had been subtracted, the average pixel value of each cell’s ROI was computed, giving F, the cell’s measured fluorescence in that frame. F0, the cell’s initial fluorescence, was defined as the average of F between the start of the experiment and the first stimulus. Each cell’s trace was then normalized as ΔF/F0, where ΔF = F - F0. All downstream analyses used this ΔF/F0 normalization, except for the PCA and learning score analyses (see below).

#### 4.4.5 Correlation-distance relationship plot

For each cell, the time-averaged (x, y) position was computed from its tracked centroid across all frames. For every pair of cells within a given culture chamber, the Pearson correlation coefficients between the ΔF/F0 fluorescence traces were computed and plotted against the pairwise Euclidean distance between mean cell positions (converted from pixels to micrometers using a calibration of 1.801 pixels/μm). Pairs were further classified by responder status into responder × responder (RR) and non-responder × non-responder (NN) categories, using the per-channel responder threshold defined above. A separate descriptive least-squares fit was drawn for the RR subset alongside the pooled fit. Because the N(N−1)/2 cell pairs are not independent (each cell appears in many pairs), the ordinary regression slope and r-value were reported as descriptive effect sizes only; statistical significance was instead assessed by a Mantel permutation test (999 permutations) per channel, which permutes cell labels, making the cell, rather than the pair, the unit of exchangeability. When pooling across channels within an experiment, per-channel Mantel r-values were combined across channels using a one-sample t-test against zero, providing a replicate-level test that treats each imaging channel (a separate chamber/cell culture; n = 3 for C2C12 cells, n = 4 for NRK-49F cells) as the replicated unit; the single-channel PC-3 dataset had n = 1 replicate, so no across-channel test was performed.

#### 4.4.6 Principal component analysis

Principal component analysis (PCA) was performed on the background-corrected fluorescence traces of all cells in each experiment, pooled across imaging channels. When channels had differing trace lengths, all channels were truncated to the common minimum number of frames. Each cell’s trace was then z-scored (mean-centered and scaled to unit standard deviation across frames), and cells with zero variance were dropped. Any residual non-finite values (e.g., from numerical edge cases) were set to zero. PCA was computed on the resulting cells × frames matrix using scikit-learn (sklearn.decomposition.PCA) with the number of components set to min(20, n_cells, n_frames). The same PCA scores were also used as input to a downstream UMAP embedding (n_neighbors = min(15, n_cells – 1); min_dist = 0.1).

#### 4.4.7 Response peak heights

For each stimulus, the response peak height was computed as the extremum of the cell’s ΔF/F0 trace within a defined post-stimulus search window, minus a pre-stimulus baseline (the mean of the five frames immediately preceding the stim), matching the baseline used for responder classification. The extremum was taken as the maximum for stimuli expected to drive fluorescence upward (DMSO pulses) and as the minimum for stimuli expected to drive it downward (acid pulses). For both DMSO perfusion experiments (C2C12 and PC3), the search window was 0.5 to 5.0 min after stimulus onset, specified in physical time and converted to an integer frame offset using each channel’s median frame interval so that the same physical window was applied despite differences in acquisition rate between cameras. For the acid feedback experiment, a frame-based window of 1 to 7 frames after the stimulus frame was used, reflecting the faster onset of the acid-driven fluorescence decrease following the 30 s acid pulse.

Per-replicate response amplitude across stimulus trains was summarized by partitioning each channel’s stimuli into consecutive trains of five pulses. Within each train, a per-cell mean response peak height was computed across the five pulses, and these per-cell train means were then averaged across all cells in the channel to yield one mean response value per (channel, train). To test for a systematic change in response amplitude across the train sequence, the difference between each channel’s last-train and first-train mean was computed, and these per-replicate differences were tested against zero with a two-sided one-sample t-test (n = number of channels), with Cohen’s d_z reported as the effect size.

#### 4.4.8 Filtering for responders

For each cell, we summarized its evoked activity with a single per-cell statistic: the mean of its per-stimulus ΔF/F0 across all N stimuli in the experiment. The per-stimulus delta was computed as the extremum of the response window minus a pre-stimulus baseline (the mean of the five frames immediately preceding the stim); using a multi-frame baseline rather than the single stim-frame value substantially reduced the tail of the noise distribution on the noisier datasets. The direction of the extremum (max for “increase” experiments, min for “decrease”) was set based on the expected cell response to each stimulus.

A null distribution for this statistic was built per (experiment, channel) by drawing N pseudo-stimuli from stimulus-free regions of the recording: frames more than max(10, response-window upper edge) away from every real stimulus, and far enough from either recording edge that both their own pre-stimulus baseline window (the first five frames are excluded) and their full response search window fit within the recording. These pseudo-stimuli were aggregated with the same mean statistic, and the procedure was repeated for 999 pseudo-replicates. The responder threshold T was the 100·(1 - α)-th percentile (α = 0.01) of |mean deltas| in the pooled null. Because the cell-level test was a single aggregate across stimuli rather than a per-stimulus test, no Bonferroni correction over the stimulus count was applied; the null SD already scales as ∼1/sqrt(N) due to averaging across N pulses. A cell was classified as a responder when its mean per-stimulus ΔF/F0 crossed the signed threshold (≥ +T for “increase”, ≤ −T for “decrease”) for that experiment, per channel. Thresholds were computed independently per channel because baseline noise levels differ.

#### 4.4.9 Habituation and sensitization scores

The habituation score quantifies the tendency of a single cell’s responses to decrease in magnitude across the stimuli within a train. For a given train of response peaks, the score was the number of times a new minimum response value was reached, not counting the first peak. The scores for each train in the experiment were summed to produce the final score. The experiments had 3 trains with 5 pulses each, so each cell’s score ranged from 0 to 12. The sensitization score was defined identically except that it counted new maxima, with the same 0 to 12 range. Both scores were computed twice per cell: once using response peak height (extremum within the response window minus the mean of the five frames immediately preceding the stimulus) and once using response width (time from the stimulus frame until the cell’s background-corrected fluorescence trace, after reaching its peak, first recrosses that same pre-stimulus baseline). Null distributions were computed by independently permuting each cell’s full 15-peak sequence across all three trains and recomputing the scores. To assess whether the observed scores could have arisen by chance, a population-level permutation test was used: the mean of the observed score distribution was compared against the means of 10,000 such shuffled distributions. The two-tailed p-value was the fraction of shuffled means whose absolute deviation from the center of the shuffled-mean distribution exceeds that of the observed mean.

#### 4.4.10 Anticipation scores

Anticipation scores measure whether single cells show unusually high or low fluorescence at the expected time of the next stimulus pulse, given the regular 10-min pulse interval used during pulse trains. The experimental protocol consists of three trains of 5 pulses each; anticipation was scored independently for the rest regions following train 1 and train 2 (train 3 had no following stimulus train). For each scored train t, the rest region was defined as the interval from the end of train t’s last response window to the start of train t+1’s first stimulus. The anticipation time was defined as 10 min after the last response peak of train t, where the peak time was taken as the across-cell median of the per-cell argmax of the background-corrected fluorescence within the response window following the train’s final pulse. For each cell, the corrected fluorescence signal across that rest region was z-scored using that cell’s own rest-region mean and standard deviation (cells with zero rest-region variance were assigned NaN and excluded). The per-cell anticipation value was the z-score at the single anticipation frame, and a per-cell shuffled control was generated by drawing, for each cell independently, one random other frame from within that train’s rest region, and recording its z-score. To test whether the cell population’s z-score at the anticipation time was significantly different from a random rest-region time, a permutation test was performed: 10,000 null distributions were generated, each by drawing one random rest-region frame per cell other than the anticipation frame, and the mean across cells of each shuffled distribution was recorded. The two-tailed p-value was defined as the fraction of shuffled means whose absolute deviation from the mean of the null distribution exceeds the absolute deviation of the observed mean from that null mean. Results were pooled across the biological-replicate channels within each DMSO experiment, excluding PC-3 as it has one replicate.

### 4.5 Molecular biology

The GCaMP6f sequence was excised from pGP-CMV-GCaMP6f (Addgene #40755; gift from Douglas Kim and the GENIE Project) [117] using BglII and NotI restriction enzymes and subsequently inserted into a pENTR1A backbone containing a CAG promoter, multiple cloning site, and SV40 polyadenylation signal (pENTR1A CAG MCS SV40PA) via the same restriction sites, generating pENTR1A CAG GCaMP6f.

For the ArcLight co. construct, the coding sequence was derived from CMV ArcLightCo (Q239)-T2A-nls-mCherry (Addgene #85806; gift from Vincent Pieribone) [118]. The insert was amplified using In-Fusion-compatible primers (forward: TTTCGAGCTCAAGCTTGCCACCATGGAAGGTTTTG; reverse: CCGCGGTACCGTCGACTCACACCTCGTTCTCGTAGC) and cloned into the pENTR1A CAG MCS SV40PA vector, which had been linearized with HindIII and SalI. Assembly was performed using the In-Fusion HD Enzyme Premix (Takara, 639649), yielding pENTR1A CAG ArcLight co.

Both entry constructs were then recombined into the piggyBac-based expression vector pmhyGENIE-3 via Gateway LR Clonase II (ThermoFisher, 11791020). This vector system is hyperactive, helper-independent, and self-inactivating, and includes a neomycin resistance cassette and was a gift from Stefan Moisyadi [119, 120]. The resulting plasmids, HypG3 NeoBB CAG GCaMP6f and HypG3 NeoBB CAG ArcLight co., were used for subsequent transfection experiments.

### 4.6 Generation of stable lines

Unmodified C2C12, PC-3, and NRK-49F cells were originally purchased from ATCC. Stable engineered cell lines were generated by transfecting cells at approximately 30% confluence with 500 ng of the relevant HypG3 plasmid using Lipofectamine 3000 (L3000008, ThermoFisher). For each well of a 24-well plate containing 500 µL of culture medium, 1 µL of Lipofectamine 3000 and 1 µL of P3000 reagent were used in accordance with the manufacturer’s protocol. After 24 hours, the transfection mixture was replaced with fresh medium, and cells were allowed to recover for an additional 24 hours prior to initiating selection with 1000 µg/mL G418. Following selection, all cells were subjected to limiting dilution in 96-well plates to isolate single clones. Individual colonies were expanded, and those demonstrating both robust proliferation and high transgene expression were selected for downstream analyses.

### 4.7 Cell cultivation

All cells were grown in FluoroBrite DMEM (Gibco A18967-01) containing 10% fetal bovine serum (ATCC), 1% penicillin-streptomycin (ATCC 30-2300, final concentrations 100 IU/ml penicillin, 100 μg/ml streptomycin), and 1% GlutaMAX (Gibco 35050-061) to make complete DMEM. Cells were propagated in 10 cm tissue culture treated dishes (Corning 430167) in humidified incubators at 37 °C and 5% CO_2_ and were passaged with trypsin-EDTA upon reaching 80% confluence.

### 4.8 Measurement of medium delay and flow pattern

When a medium containing a drug is loaded into a reservoir of the CellASIC® ONIX2 plate, the fluidic channel between it and the cell culture chamber initially contains medium without the drug, and the drug reservoir must be perfused for some length of time before it flushes all the way through the channel and reaches the cell chamber. After this point, when the drug reservoir is perfused again, drug reaches the cell almost instantly. To measure this “initial delay time,” a technique similar to residence time distribution measurement was used [88]. The reservoirs of a CellASIC® ONIX2 chamber were loaded with deionized water and perfused to fill the channels and chamber with water. A tracer fluid was made by dipping a yellow highlighter pen into water, making a fluid that fluoresces with blue excitation. The tracer was loaded into reservoirs 2 and 5, the same ones used for the drug in subsequent experiments with DMSO. Perfusion of the tracer reservoirs was started at 2.5 psi (the same pressure used in subsequent experiments), and simultaneously, fluorescence imaging of the chamber was started, imaging every 7 seconds. Two trials showed that the dye was first detectable in the images after 89.5 seconds on average. This was rounded to 90 seconds (1.5 min) and used as the initial delay time in subsequent experiments. The perfusion and imaging were continued until the entire chamber was filled with tracer in order to observe the flow pattern of the incoming tracer as it replaced the water in the chamber. This revealed a complex pattern in which the tracer initially flowed in along the walls of the chamber before filling the center.

### 4.9 Microfluidic plate preparation

The cell chambers of the CellASIC® ONIX2 microfluidic solution switching plates for mammalian cells (M04S) were pre-coated according to manufacturer instructions with fibronectin solution (15 μl of fibronectin solution (1 mg/ml, Sigma-Aldrich F1141) per 1 ml FluoroBrite DMEM without additives, giving 0.015 mg/ml). A suspension of 1×10^4^ cells/ml was prepared in complete DMEM and each chamber was seeded with a 10 μl droplet, according to manufacturer instructions. Chambers were perfused by a gravity-driven of complete DMEM and were used in experiments the following day.

Perfusion medium was then prepared, consisting of FluoroBrite DMEM with 2% FBS to reduce background fluorescence, with 1% penicillin-streptomycin and 1% GlutaMAX. The medium was exposed to blue LED light (470 nm) for 30 min to pre-photobleach fluorescent medium components to prevent dynamics in background fluorescence from occurring during experiments. The perfusion medium was kept overnight in the cell culture incubator in a 10 cm dish to pre-condition its temperature and CO_2_ content.

The microfluidics of the CellASIC® ONIX2 plates come pre-filled with phosphate buffered saline (PBS). Immediately before experiments, perfusion medium (without drugs) was loaded into the plate’s reservoir wells, all of which were then perfused at 2.5 psi for 5 min to flush the PBS out of the fluidics and prime them with medium.

### 4.10 C2C12 DMSO feedforward experiment

Pre-conditioned perfusion medium (normal medium, N) was loaded into reservoirs 3 and 4 for each of 3 cell chambers of the CellASIC® ONIX2 plate. Preconditioned medium with 5% DMSO (50 ul DMSO/ml of medium, Sigma-Aldrich D2438) was prepared (DMSO medium, D) and immediately loaded into reservoirs 2 and 5 for each chamber.

One imaging position was set for each chamber and the imaging loop was started, with each location being imaged every 21 seconds. Simultaneously, the CellASIC® ONIX2 perfusion program was started, perfusing N and D medium as follows (for N, reservoirs 3 and 4 were perfused at 2.5 psi, while for D, reservoirs 2 and 5 were perfused at 2.5 psi): 13.5 min N, 1.5 min D (measurement of the “initial delay time” above found that it takes 1.5 min for the DMSO medium to travel from the reservoirs to the chambers the first time it is dispensed, resulting in the cells experiencing N for the first 15 min of the experiment), cycle 5x (2 min D, 8 min N), 30 min N, cycle 5x (2 min D, 8 min N), 30 min N, cycle 5x (2 min D, 8 min N).

### 4.11 PC-3 DMSO feedforward experiment

The PC-3 DMSO experiment was conducted in the same manner as the C2C12 DMSO experiment, except 1 cell chamber was used and was imaged every 7 seconds, and the cells were given a total initial perfusion acclimation time of 10 min instead of 15 min.

### 4.12 NRK-49F acid feedback experiment

Acidic medium (A) was prepared by adding 15 μl of 1 N HCl per ml of perfusion medium and allowing it to equilibrate in a CO_2_ incubator for at least 2 h, resulting in a pH of 1-2 as measured with MQuant pH indicator strips. This amount of acid was chosen because cells could tolerate it in short exposures and it gave a strong fluorescence change signal. Perfusion medium (N, normal medium) without acid had a measured pH of 7-8.

In our setup, two CellASIC® ONIX2 chambers received simultaneous independent control. Each chamber could be in 1 of 2 states at any time: perfusing N, or perfusing A. The plate has an array of medium reservoirs (4 reservoirs per chamber); each row supplies one chamber, and whole columns must be dispensed at the same time. Thus, to cover all possible states, the reservoirs for the first chamber were filled with A, A, N, and N medium (reservoirs 1-4), while the second chamber’s reservoirs were filled with A, N, A, and N. Any possible perfusion state of the two chambers could then be covered by activating one column of reservoirs (AA, AN, NA, NN).

One imaging position was set for each chamber, and an initial fluorescence image was taken of each. These images were then segmented to locate the cells. Each chamber was imaged again during a manually delivered 30 second pulse of A to determine the dynamic range of the population average ArcLight fluorescence. A setpoint fluorescence level was chosen for each chamber between the levels of these two images, approximately 1 fluorescence unit below the initial value. The experiment was started with an imaging interval of 20 seconds and a constant perfusion of N at 2.5 psi to each chamber for 5 minutes with the controller off.

After 5 minutes, the controller python script was turned on. Whenever a new set of cell images for the two chambers was saved to the directory, the controller computed the mean cell fluorescence from the raw pixels using a cell segmentation mask defined prior to starting the experiment. The control policy was a simple If statement. The device delivers normal perfusion medium by default, but if the average fluorescence of the latest image was greater than a pre-defined threshold, then a 30 second pulse of acidic medium is delivered. After each new image, the correct perfusion state for the two chambers was determined (AA, AN, NA, or NN), and the correct reservoir column was activated. The controller sent commands to the CellASIC® ONIX2 via its API to stop the current experiment file and start up the file with the correct state.

## Supporting information

NRK-49F Movie A

NRK-49F Movie B

NRK-49F Movie C

NRK-49F Movie D

Hardware Schematic - LED ring diagram

Hardware Schematic - LevinScope

SLDDRW and STEP file of LevinScope Hardware Schematic

## Acknowledgements

We thank Ann Karim, Navneet Jawanda, and Robert Brucker for project management support, Rakela Colon for laboratory management, and Tomika Gotch for assistance with manuscript preparation, along with Tom Cirrito, Suomia Abuirqeba, and many members of the Levin lab for helpful technical discussions. The authors acknowledge the Tufts University High Performance Compute Cluster https://it.tufts.edu/high-performance-computing.

## Funding

M.L. gratefully acknowledges support of Templeton World Charity Foundation Inc., John Templeton Foundation grant 62212, and Astonishing Labs. The opinions expressed in this publication are those of the authors and do not necessarily reflect the views of the sponsors.

## Conflicts of interest

This research was partially funded at Tufts under a Sponsored Research Agreement with Astonishing Labs. M.L. is a co-founder and shareholder of Astonishing Labs. Astonishing Labs has certain rights to any inventions associated with this research.

## Supplemental Materials

The Supplemental Materials of this manuscript include high-resolution schematics for the optical system and movies of all cell chambers (downsampled to 10% of their original sizes, but otherwise unedited). The code used for this work is publicly available under a modified Apache license in two GitHub repositories. The code for operating the custom mobile microscope can be found here https://github.com/k-shcherb/levin_lib/tree/Publish. The code for running closed-loop experiments (analyzing images in real time and sending commands to the CellASIC® ONIX2 through its API), and for all image and data analyses can be found here https://github.com/douglashazel/closed-loop-cell-imaging. This repository also contains the segmentation masks, raw fluorescence traces, and normalized fluorescence traces, for all cell chambers in this paper.

## Notes

https://github.com/k-shcherb/levin_lib/tree/Publish

https://github.com/douglashazel/closed-loop-cell-imaging

