## Supplementary figures and images for "A platform for automated training of mammalian cell physiology"

### Hardware Schematic - LED ring diagram

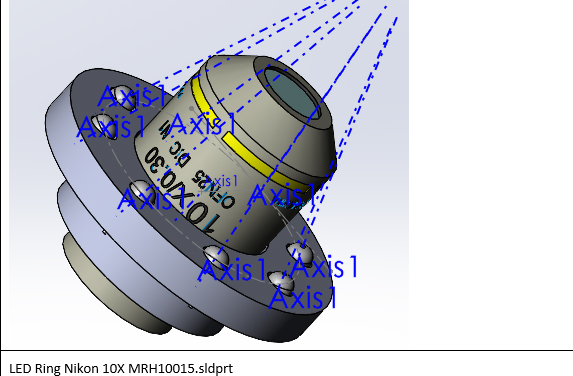
